# Genome-wide SNP detection in Darjeeling tea: unravelling genetic structure, functional impact and trait associations

**DOI:** 10.1101/2020.09.02.280461

**Authors:** Anjan Hazra, Rakesh Kumar, Chandan Sengupta, Sauren Das

## Abstract

Genotyping by sequencing and identification of functionally relevant nucleotide variations in crop accessions are the key steps to unravel genetic control of desirable traits. In the scope of present work, elite cultivars of Darjeeling tea were undergone SNP genotyping by double-digest restriction site associated DNA sequencing method. This study reports a set of 54,206 high-quality SNP markers discovered from ∼10.4 GB sequence data, encompassing 15 chromosomes of reference tea genome. Genetic relatedness among the accessions conforms to the analyses of Bayesian clustering, UPGMA, and PCoA methods. High percent of heterozygous loci in the majority of the cultivars reflect their ‘hybrid’ ancestry as reported earlier. Genomic positions of the discovered SNPs and their putative effect on annotated genes designated a thoughtful understanding of their functional aspects in tea system biology. A group of 95 genes was identified to be affected by high impact variants, most of them are involved in signal transduction, biosynthesis of secondary metabolite, transcriptional and translational regulation. Genome-wide association analyses of 21 agronomic and biochemical phenotypes resulted in trait-linked polymorphic loci with strong confidence (*p* < 0.05 and 0.001). The selection of significant marker-trait associations with the Bonferroni correction threshold retained a set of 57 SNPs distributed across 14 chromosomes to be linked with eight phenotypic traits. High impact and trait-associated nucleotide polymorphisms perceived in this study can be exploited in worldwide germplasm of contrasting origin to depict their heritability and to unlock their potentiality in marker-assisted breeding.

## Introduction

Tea, the globally second most-consumed drink after water, is popular for its irreplaceable refreshing value and potential health benefits. Following industrial manufacturing of the fresh leaves from perennial evergreen *Camellia sinensis* plants, processed tea possesses a certain distinctive set of secondary metabolites that contribute to its pleasant aroma, taste, and other wellbeing attributes (Hazra et al., 2019; Xia et al., 2020). Although the plant itself grows in diverse habitats, the coziness and quality vary with the geographical location and cultivar origin (Firmani et al., 2019; Hazra et al., 2018a). Since its inception, Darjeeling tea, in particular, is well known for its premium quality and brand value in the world beverage market. The temperature-moisture combination of Darjeeling hills prevailing throughout the year, the genetic background of the planted bushes, and overall the undergoing processing stages have been ascribed to the generation of a majestic flavor associated with the superior quality of this tea (Das, 2006). Owing to the excellence and physiognomies, Darjeeling tea as the first Indian product, achieved Geographical Indications tag in 2004–2005 (Das, 2006) so that the cultivation and production of the same are maintained and restricted to the geographical territory of Darjeeling hills. The report states that the production of Darjeeling tea contributes only 1% of the total Indian tea production, however, in terms of economic paybacks, it supersedes all others since last considerable period (Hazra et al., 2018b). Nonetheless, the recent gradual changes in temperature and rainfall patterns in Darjeeling hills imposing an alarming threat to the legendary Darjeeling tea industry leading to deterioration in overall production (Singh, 2019).

Being a perennial crop and evolved to thrive in diverse ecological conditions, specific cultivars of tea plants with an anticipated genetic makeup, possess unique traits of climate-smart attributes. In addition to various stress resilience, they can also serve as an efficient and sustainable natural exhauster by capturing and storing CO_2_ in their dense green coverage along mountain slopes and valleys (Phukan et al., 2018). Hence, the selection and utilization of vigor cultivars with desirable phenotypes are crucial toward the prospective crop improvement program in tea. Development of trait-associated molecular markers has been adopted for tea in the recent past (Hazra et al., 2018a; Mukhopadhyay et al., 2016). Due to several bottlenecks in the life cycle, conventional association mapping has been found difficult in tea (Hazra et al., 2020a). Hence, simultaneous genotyping and phenotyping of contrasting core accessions and subsequent marker-trait association studies were suggested as a convenient choice to overcome the challenges (Hazra et al., 2019; Hazra et al., 2018a).

Single nucleotide polymorphisms (SNPs) are considered as a superior option among the molecular markers because of their high frequency in genomes and expedient detectability using high-throughput approaches (Guajardo et al., 2020; Mammadov et al., 2012). SNPs can be accepted as key factors for the generation of phenotypic variation and their effect on functional alteration of genes enlightens the functional genomics of an organism (Hirakawa et al., 2013). Genotyping-by-Sequencing (GBS) applies next-generation sequencing approaches to enable both the *de-novo* and reference-based genotyping of numerous SNPs in a diverse set of samples (Elshire et al., 2011). Among the several types of high-throughput SNP genotyping methods discovered in recent years, double-digest restriction site associated DNA (ddRAD) sequencing is one of the most convenient, cost-effective, and widely used platforms (Peterson et al., 2012; Ray and Satya, 2014). This genome-wide approach consecutively involves (i) low- and high-frequency cuts to digest the DNA, (ii) ligation of a barcoded adapter to one restriction site and a common adapter to the other, (iii) pooling, (iv) size selection, (v) library enrichment, and (vi) introduction of a second barcode in the form of an Illumina index to increase multiplexing (Scheben et al., 2017). The ddRAD generated sequence data have been employed for various plant breeding approaches like linkage mapping (Davik et al., 2015), QTL analyses (Chen et al., 2017; Laila et al., 2019; Sasai et al., 2019), and genome-wide association studies (Jaiswal et al., 2019; Luo et al., 2019; Raggi et al., 2019; Sudan et al., 2019). Assessment of the genetic diversity of different plant species, such as apple (Ma et al., 2017), orchid (Roy et al., 2017), *Prunus* (Guajardo et al., 2020), Sesame (Basak et al., 2019) and onion (Lee et al., 2018) have also been carried out through ddRAD technology. In tea, ddRAD based SNP genotyping was applied for the determination of genetic diversity and population structure among the germplasm accessions and their wild relatives from China and Japan (Yamashita et al., 2019; Yang et al., 2016).

Darjeeling tea cultivars of diverse genetic makeup that carrying contrasting phenotypic aptitudes are being maintained at the experimental field of Darjeeling Tea Research and Development Centre for research and trial purposes. Dissecting the genetic background of the desirable phenotypes underlying these accessions is needed to excavate their breeding potential. Extensive phenotyping and simultaneous genotyping are considered the most suitable option for attending the resolution of efficient breeding (Raggi et al., 2019). Sequence-based genetic characterizations, especially with SNP markers are too inadequate to uncover the potential of these genomic resources in India. To the best of our knowledge, ddRAD based, or any other GBS for Darjeeling tea cultivars have not been reported so far. In this study, the core collection of 23 elite accessions for the cultivation of Darjeeling tea were taken into account for the ddRAD-seq assisted genotyping approach. Identified SNPs distributed throughout the tea genome were then assessed to derive the levels of molecular diversity and genetic structure among the accessions, extrapolation of SNP effects, and identification of SNP-carrying genes. Genome-wide association studies with the high confidence variant sites were accompanied for the major agronomic traits of tea, to harvest desirable trait-associated loci. SNPs with potential functional impact and/or phenotypic trait relations were presented for possible implementation in the marker-assisted selection and thereby forthcoming crop improvement programs of tea.

## Materials and Methods

### Plant materials

A total of twenty-three elite tea cultivars released for plantation at the various slopes of Darjeeling hills were chosen for this study. All these accessions are being maintained at Darjeeling Tea Research and Development Center (DTRDC), Kurseong, with uniform agro-ecological conditions for research and breeding purposes (Hazra et al., 2020a; Hazra et al., 2020b). Most of the cultivars are commercial hybrid, generated through preferential breeding techniques among the wild and domesticated types of tea lineages. Uniqueness in yield and flavor potential of the commercial accessions were described earlier, through evaluation in long-term field trial (Singh, 1992). These include the widely cultivated varieties by Darjeeling tea planters for their excellent aroma traits (AV-2, K-1/1, and B-668), very high yield (HV-39 and T-78), or drought-tolerant characteristics (RR/17/144). Fresh leaf and bud samples were harvested from each of the studied cultivars, kept in frozen condition, transferred to the laboratory, and stored in −80°C refrigerator until further use.

### Genotyping by sequencing and computational processing

Genomic DNA extraction from harvested young leaves of each of the cultivars was performed using the DNeasy Plant Mini Kit (Qiagen) according to the manufacturer’s instructions. The quality and quantity of the isolated DNA were examined by 1% agarose gels (Figure S1) and a Qubit Fluorometer v2.0 (Thermo Scientific, USA) respectively. For GBS, a double digest restriction associated DNA (ddRAD) sequencing protocol was employed. A total of 1 µg of intact genomic DNA for each sample was digested using *Sphl* and *Mluc1* restriction enzymes. Digested products were cleaned up using Ampure XP beads (Beckman Coulter Genomics) followed by ligation of P1 (barcoded) and P2 adaptors using T4 DNA ligase. Following subsequent pooling and cleanup, the size selection of the ligated products was carried out by 2% agarose gel electrophoresis. PCR amplification was done to enrich and add the Illumina specific adapters and flow cell annealing sequences. QC of the prepared ddRAD libraries for each of the samples was checked using a Bio Analyser (Agilent Technologies, USA). All the 23 ddRAD libraries with satisfactory QC profiles were then sequenced on an Illumina HiSeqX 10 platform (AgriGenome Labs Pvt. Ltd., India) using a 2×150 bp paired-end read module. Raw sequence reads were demultiplexed and filtered based on RAD tags at both ends of the pair. The quality of paired-end raw reads was examined in FastQC (Andrews, 2010). Filtered reads were pre-processed in Trimmomatic-0.33 (Bolger et al., 2014) for removal of the Illumina adapters and 5′ and 3′ base trimming. Subsequently, the remaining clean reads were subjected to paired-end alignment against the tea plant reference genome (Xia et al., 2020) using Bowtie v2.3.4.3 (Langmead et al., 2009) with default parameters. The ddRAD sequencing generated raw reads has been deposited in the NCBI Sequence-Read Archive (SRA) database.

### Genome-wide SNP detection

Reference genome aligned sample-specific BAM files were sorted by coordinate, de-duplicated, and assigned appropriate read groups with the help of SAMtools (Li et al., 2009) and Picard Toolkit (http://broadinstitute.github.io/picard/). Based on the preprocessed BAM files, the SNP calling was performed using FreeBayes (Galaxy Version 1.3.1) (Garrison and Marth, 2012) on a galaxy interface (Afgan et al., 2018). The bi-allelic SNPs were filtered at minimum read depth 10 and de-complexed using the VCFlib toolkit (Garrison and Marth, 2012). Identified polymorphic SNPs with 10X read coverage were further subjected to filtering for minor allele frequency (MAF) ≥ 0.05.

### Estimation of SNP stats, heterozygosity and population structure

SNP densities across 15 chromosomes of the tea plant were represented by histogram and heatmap using Circos program v0.69-8 (Krzywinski et al., 2009) and plotted in 1 Mbp window size. Minor allele frequency and Tajima’s D distribution of the polymorphic sites, frequencies of SNP across chromosomes, and transition-transversion ratio were estimated via the SNIplay v3.0 tool (Dereeper et al., 2015). Determination of heterozygosity percent, UPGMA clustering, and PCoA analyses of the accessions were carried out in TASSEL v5.2.64 (Bradbury et al., 2007). To elucidate the genetic structure among cultivars, a Bayesian clustering algorithm approach was employed via the admixture model with correlated allele frequencies in STRUCTURE v2.3.4 (Pritchard et al., 2000). Before this, the SNP containing (.vcf) file was converted to the corresponding software input file formats by PGD Spider v2.1.1.5 (Lischer and Excoffier, 2012). A total of 10 independent runs were evaluated for 1 to 5 genetic subgroups, which were assumed to infer the optimum number of populations (K) present within the accessions. For each run, a burn-in period of 50000 and a Markov Chain Monte Carlo replication of 100000 were set. Most probable K (the number of subpopulations) was determined with the ΔK statistics via Structure Harvester v0.6.94 (Earl, 2012). The bar plots of population structure were generated with genotype labels and ordered by Q-value.

### Functional effects of SNPs on genome

Identified high confidence SNPs among Darjeeling tea cultivars were mapped to the chromosome level reference genome of tea (Xia et al., 2020) to ascertain the putative candidate genes residing at their close vicinity. Associated SNPs effects on the genome were predicted using SnpEff v4.3 (Cingolani et al., 2012) based on a locally created database with the gene annotation (gff3) file retrieved from TPIA (Xia et al., 2019). Transcripts encompassing the variants within an exon, intron, splice site, and 5 Kbp up- or down-stream positions were recorded along with their description. The occurrence of SNP and their predicted effects were subsequently categorized as high (disruptive impact on the encoded protein), moderate (non-synonymous substitution), low (synonymous substitution), and modifier (impact on noncoding regions) impacts. Significant SNP associated genes were grouped according to their functions based on the KOG annotation database.

### Marker trait association

Collectively 21 tea specific traits included in the present investigation as reported agronomic characters (yield, flavor, and stress tolerance) (Bhuyan et al., 2012; Singh, 1992) and our earlier findings on rigorous phenotyping of the cultivars (biochemical traits, Hazra et al. (2020a); stomatal characters, unpublished), were considered for possible marker-trait association studies. Accurately filtered 54,206 SNPs from 23 genotypes were considered for the GWAS. Genome-wide marker-trait association (MTA) analyses were conducted using TASSEL v5.2.64 software (Bradbury et al., 2007) with the GLM protocol keeping a significant threshold for the association at *p* < 0.05 and *p* < 0.001. Furthermore, the Bonferroni equation derived *p*-value has been fixed as a threshold for the significance of putative associations between the SNPs and corresponding traits of interest. For the model fitting test of each trait, quantile-quantile (Q-Q) plots were generated showing the distribution of expected and observed *p*-values from the association test output. The Manhattan and QQ plots were represented using the ‘qqman’ R package (Turner, 2014) displaying the significant trait-associated markers.

## Result

### Genotyping

The present study acquired a total of 69,540,256 raw reads through the ddRAD based paired-end (150×2) sequencing on an Illumina Hiseq X10 platform. This output spanned an average of 3.02 M reads and 43.45 % GC content per accession with the highest (5.73 M) and lowest (1.46 M) read observed for the cultivars Badamtam-15/263 and Teesta Valley-1 respectively (Table S1). Pre-processed reads with RAD tag at both loci (98.44 ± 0.5 in percent) were mapped to the tea plant (*Camellia sinensis var. sinensis*) reference genome and resulted in a mean of 90.5% alignment rate (Table S1). Initially, 2,120,435 SNPs were identified through the variant calling pipeline, which on screening with 10X read depth and Q30 quality score cut-off, retained a total high quality 2,24,966 SNPs (Table S2).

Nucleotide substitution pattern of the variants included 1,65,567 transitions (Ts) and 59,399 transversions (Tv), with the transitions to transversions ratio of 2.79. The incidences of various substitutions were 36.9% C/T, 36.7% A/G, 6.2% A/T, 7.3% A/C, 7.4% G/T, and 5.5% C/G (Figure 1A). Tajima’s D and Minor allele frequency distributions of the polymorphic sites indicated that a predominant amount of positive Tajima’s D value and abundant sites with low MAF were present (Figure 1B-C). SNPs mapped to the chromosomes ranged from an average 82.8 (Chr3) to 121.6 (Chr1) per megabase (Table 1, Figure 1D). Variant sites residing in 15 chromosomes were further filtered with stringent parameters (minor allele frequency ≥ 0.05, missing data within a locus < 0.3) for high confidence marker-trait association analyses, which returned a total of 54,206 SNPs.

**Table 1.** Distribution of identified SNPs mapped to each chromosome.

| **Chromosome** | **Length in bp** | **No. of genes residing** | **Detected SNPs** | **Mean SNPs per Mb** |
| --- | --- | --- | --- | --- |
| Chr1 | 2.23E+08 | 3850 | 27106 | 121.6 |
| Chr2 | 2.13E+08 | 3814 | 22660 | 106.3 |
| Chr3 | 1.87E+08 | 3258 | 15519 | 82.8 |
| Chr4 | 1.96E+08 | 3314 | 19684 | 100.3 |
| Chr5 | 1.95E+08 | 2843 | 18283 | 93.8 |
| Chr6 | 1.81E+08 | 3332 | 16745 | 92.4 |
| Chr7 | 1.87E+08 | 3301 | 18884 | 100.9 |
| Chr8 | 1.63E+08 | 2237 | 18547 | 113.7 |
| Chr9 | 1.66E+08 | 2737 | 17884 | 108.0 |
| Chr10 | 1.67E+08 | 2492 | 15982 | 95.6 |
| Chr11 | 1.24E+08 | 2374 | 13808 | 111.6 |
| Chr12 | 1.63E+08 | 2332 | 15626 | 96.0 |
| Chr13 | 1.36E+08 | 2345 | 13209 | 97.0 |
| Chr14 | 1.34E+08 | 2367 | 13771 | 102.9 |
| Chr15 | 1.19E+08 | 2009 | 11929 | 100.2 |
| Unassigned Contigs | 2.99E+08 | 7920 | 20971 | 70.1 |

**Figure 1.**
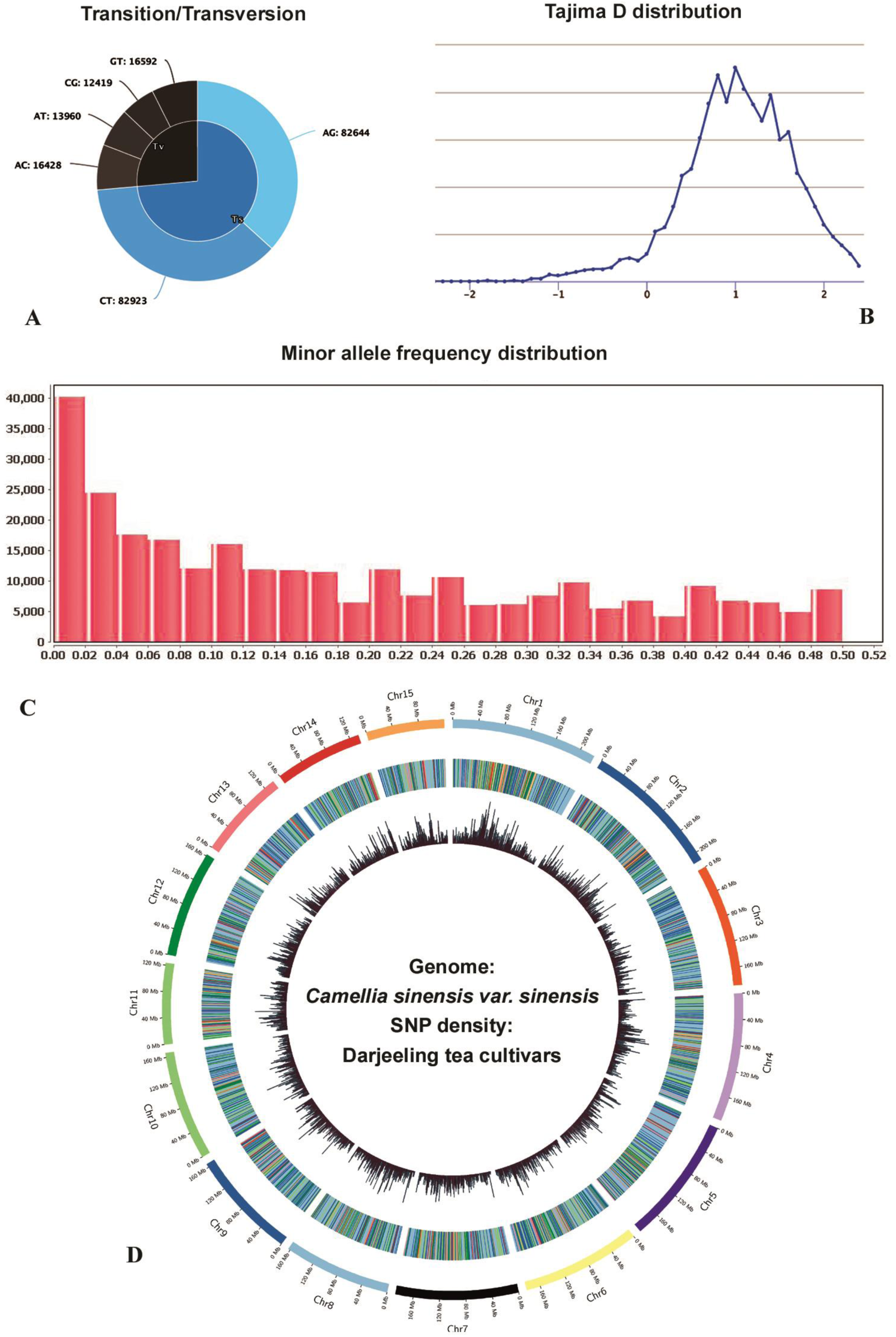
Characteristics of genome-wide SNPs identified. A – Transition and transversion fractions, B – Tajima’s D distribution of identified variant sites, C – minor allele frequency distribution of variant sites with 10X read depth, D – Distribution of SNPs along the 15 chromosomes of tea genome (*Camellia sinensis var. sinensis*). SNP density was plotted in 1 Mbp sliding window using Circos. Heatmap and histogram tracks from outside to inside represents distribution of initial high depth (10X) 2,24,966 SNPs and subsequently filtered (MAF < 0.05, minimum SNP call in a locus > 0.7) 54,206 SNPs respectively.

### Varietal diversity and genetic structure

Genome-wide SNPs were utilized to clarify the genetic structure among the studied elite cultivars of Darjeeling tea. The optimal number of subgroups (K) determined by STRUCTURE software run and subsequent Structure Harvester evaluation suggested peak ΔK value at K = 2 followed by K = 3 (Figure 2A). Bar plots of structure analyses considering K = 2, grouped the cultivars like MB-6, P-312, B-777, AV-2, and S-1 in a single subpopulation, TS-569 and RR-17/144 to be admixed and the remaining accessions classified to another set (Figure 2B). When ancestral components are assumed K = 3, the formerly mentioned group were intact as a single population, however, the larger part of accessions displayed admixture between two other subgroups at varying degrees. In the same output, cultivars like TS-569 showed a membership of all three subgroups (Figure 2C).

**Figure 2.**
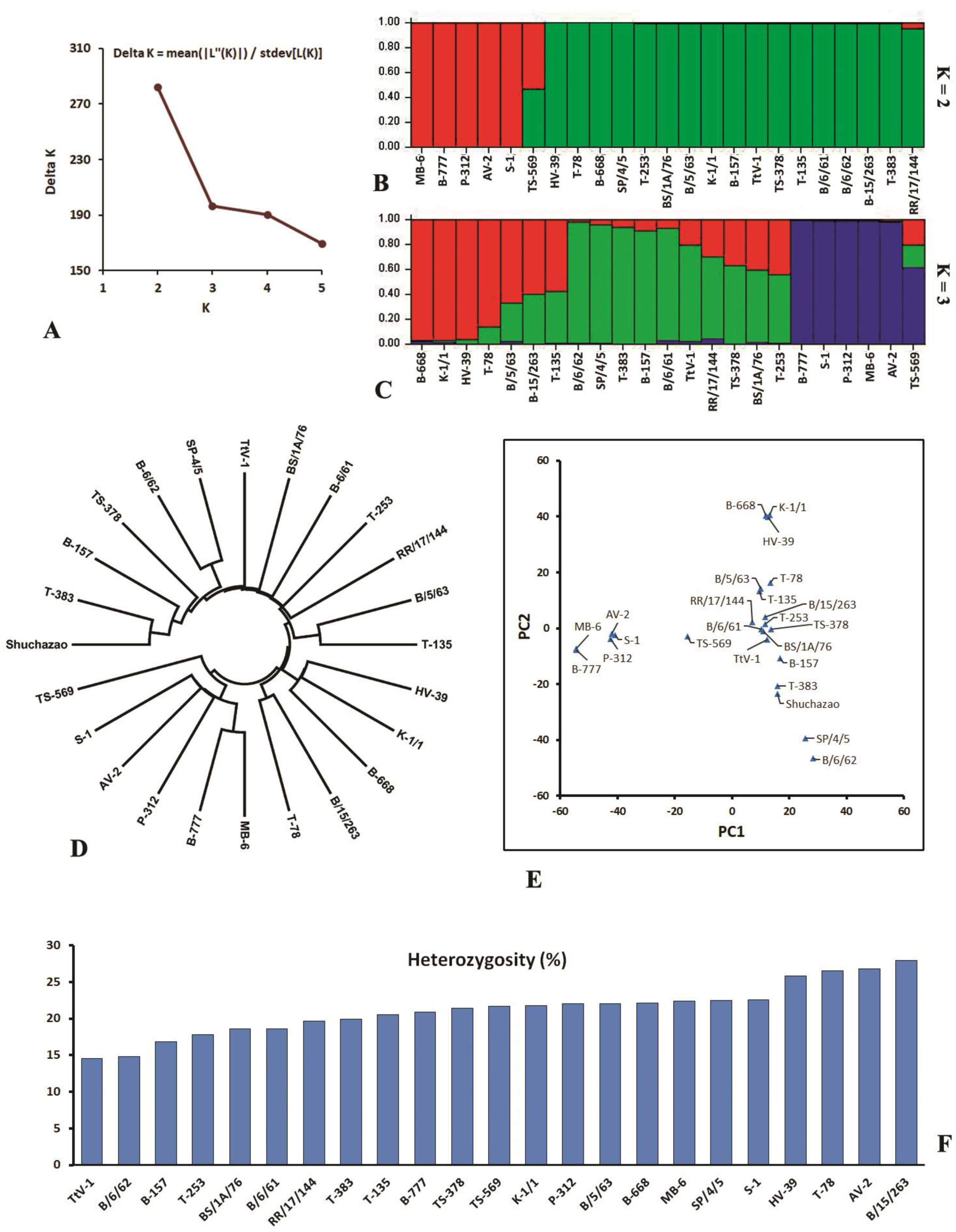
Genetic relatedness and heterozygosity levels of 23 Darjeeling tea cultivars. A – Optimal number of subpopulations estimated by plotting ΔK against each K of STRUCTURE analyses, B – Inferred genetic structure at K = 2 and C - K = 3, D – UPGMA dendrogram and E – Principal coordinate analyses with the identified SNP markers, F - Heterozygosity percentages of the tea accessions.

UPGMA dendrogram analysis has clustered the accessions in two distinct subgroups (Figure 2D). One of them was grouped by MB-6, B-777, P-312, AV-2, S-1, and TS-569 being separated from the main subgroup. In other subgroups, few cultivars strongly clustered with each other suggesting their genetic similarity, such as T-135 and B/5/63 belonged to the same branch. Likewise, two other cultivar pairs - B/6/62, SP/4/5, and T-78, B/15/263 shared a single subcluster of each. Results of the principal coordinate analyses were strongly corroborating to the genetic structure and dendrogram analyses (Figure 2E). Particular coordinate held by an accession and occurrence of others in the vicinity reflected the major subgroups as revealed by K = 2 and k = 3 in structure results as well as sub-clustering patterns of each cultivar as observed in the tree. Percentage of heterozygosity within the accessions indicated the lowest amount of heterozygous sites in TtV-1 (14.54) and B/6/62 (14.76), while the maximum value occurred in B/15/263 (27.94), AV-2 (26.75), T-78 (26.5), and HV39 (25.8). Heterozygosity of the remaining cultivars are almost similar and lies between 17-22% (Figure 2F).

### SNP effects on genome

Prediction of 54,206 well-confidenced SNPs putative effect on tea genome exhibited most of them having modifier effect (96.06%), followed by moderate effect (2.1%). Least occurrence was observed in case of high impact (0.22%) *i*.*e*. with disruptive impact on the encoded protein. SNPs with modifier impact were frequently observed in non-coding genomic regions, which are intergenic positions (75.96%) followed by intron variants (8.33%) and within 5kb downstream (6.86%) or upstream (4.22%) of the annotated genes. The present study revealed a total of 3177 genes in the tea genome to be impacted by SNPs at varying degrees. According to the functional impact classification of the SNPs in the exon region, missense type was the most prevalent one (59.79%) and a little expanse (1.79%) of nonsense mutations were observed. A considerable amount of variants occurred in acceptor or donor sites of splicing regions too (Figure 3, Table S3).

**Figure 3.**
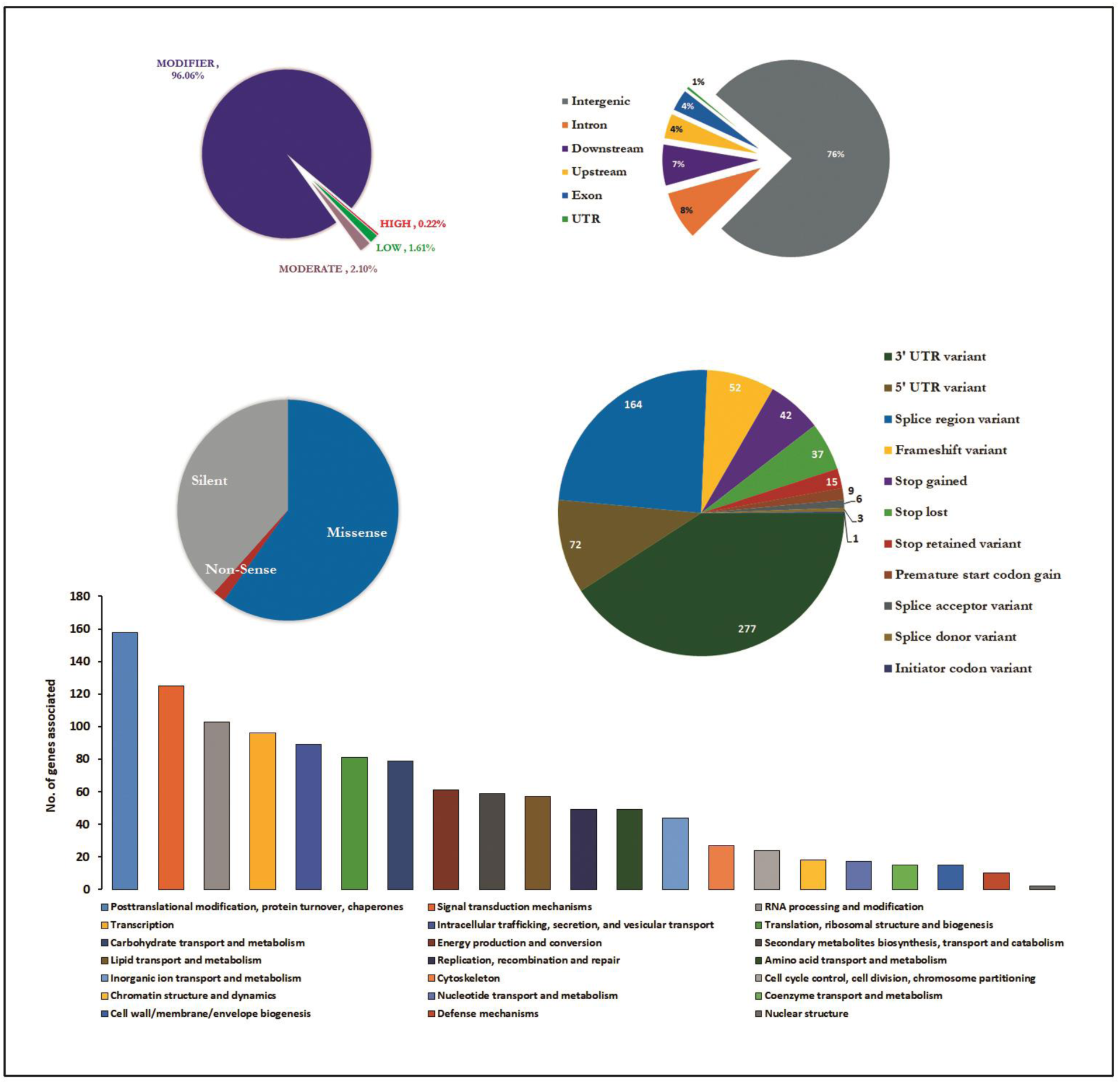
Classification of detected SNPs according to their effect and distribution across various genomic regions (Pie charts). The bar graph represents the KOG annotation-based classification of genes that harboring variant sites with potential effects.

A total of 95 genes were identified to harbor variant sites that have a potential impact on the gene function by altering their protein product and thereby probable phenotype modification of the source plant (Table S4). Most of these genes were with at least one SNP except for CSS0030092 which had four and CSS0004975, CSS0008917, CSS0011515, CSS0017046, CSS0019151, CSS0029404 with three in each, and 28 genes having two SNPs. The KOG class prediction ensued that a major fraction of SNPs impacted genes are involved in posttranslational modification, signal transduction mechanisms, RNA processing, and transcript modifications, etc (Figure 3). However, 46.74% of the candidate genes found no hit to the KOG database, and 16.18% of them were poorly characterized (*i*.*e*. General function prediction only or unknown function) (Table S5). Predicted changes on encoded amino acids from variant coding sites led to various modes of consequences at the protein level. These include nonpolar to polar (Ala-Thr, Gly-Arg, Pro-Arg, etc.), negatively charged to positively charged or non-charged (Glu-Lys, Asp-Asn, etc.) and aromatic to aliphatic (Phe-Leu) residue substitutions (Table S6) as a major fraction of occurrences.

### Genome-wide association studies

The core set of high quality (MAF ≥ 0.05, missing data < 0.3) SNPs were chosen for association analyses with 21 agronomic and health benefit traits that are specific to tea. The probability of association between each trait and putatively associated variant sites was depicted using Manhattan and QQ plots (Figure 4a, b). A certain amount of loci were observed to show covariance with the phenotypic traits at varying probability threshold (*p* < 0.05 and *p* < 0.001) (Table S7). However, to eliminate the low confidence associations and reduce false-positive rates, a stringent screening was performed with the manual examination of expected and observed probabilistic QQ plots followed by the selection of putatively associated SNPs by a Bonferroni correction threshold (*p* < 1.845 e^-05^). Accordingly, a total of 57 markers were identified to be significantly associated with eight phenotypic traits (Table 2). These trait-associated markers were distributed across the 14 out of 15 chromosomes of tea. Most of the significant associations observed in case of various secondary metabolites, such as EGCG and flavonoids. Mapping of the SNPs on the genome resulted in a major amount of the markers localized on intergenic regions. However, a few were observed to occur in the downstream, upstream, or intron portions of the gene as well. One flavonoid related SNP was observed to cause a missense type of mutation in the CSS0000649 gene. KOG class prediction of the trait-associated SNP impacted genes resulted in signal transduction related genes is the most predominant followed by the post-transcriptional and post-translational type.

**Table 2.**
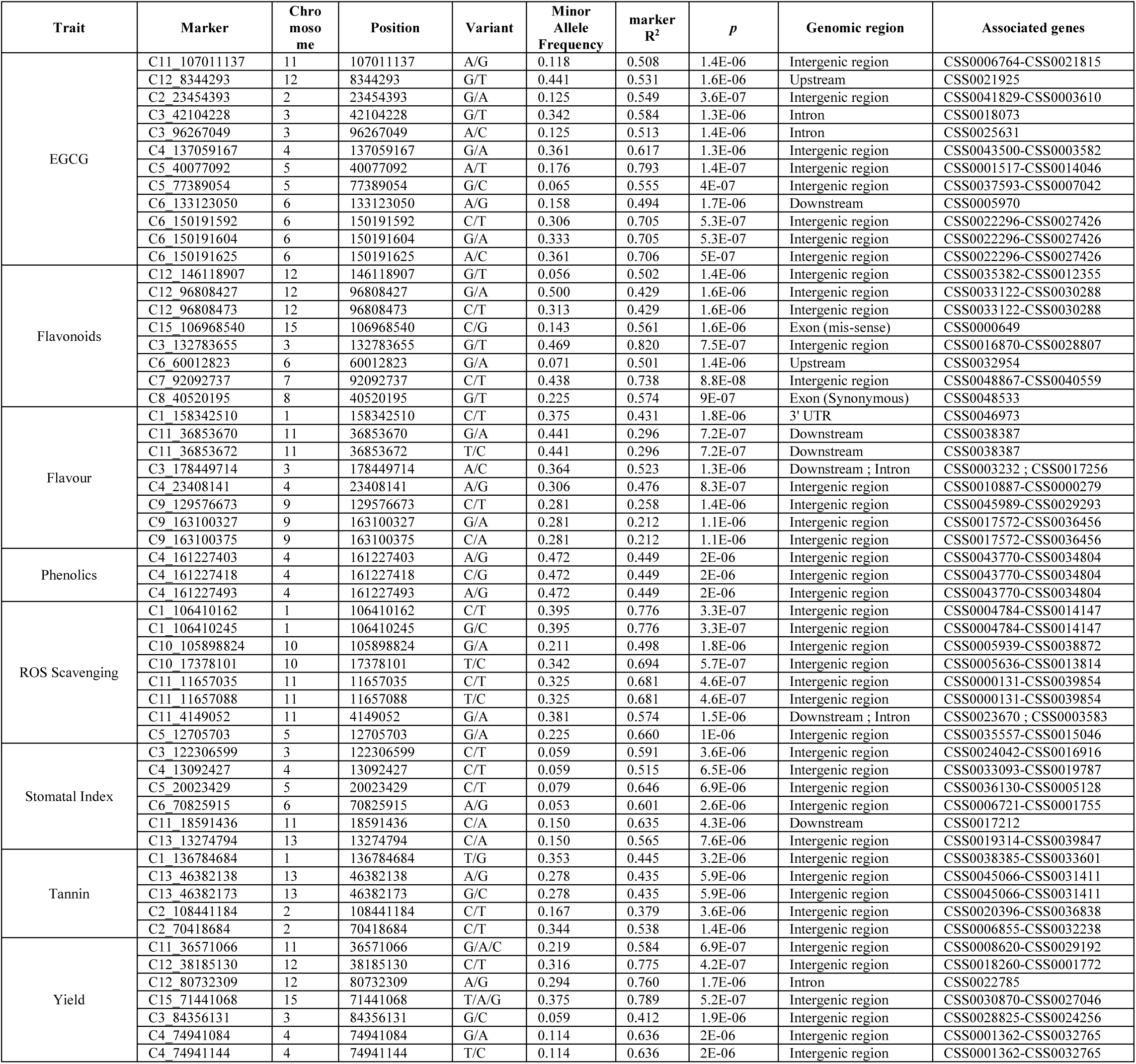
Summary of trait-associated markers that accomplished the Bonferroni correction threshold.

**Figure 4.**
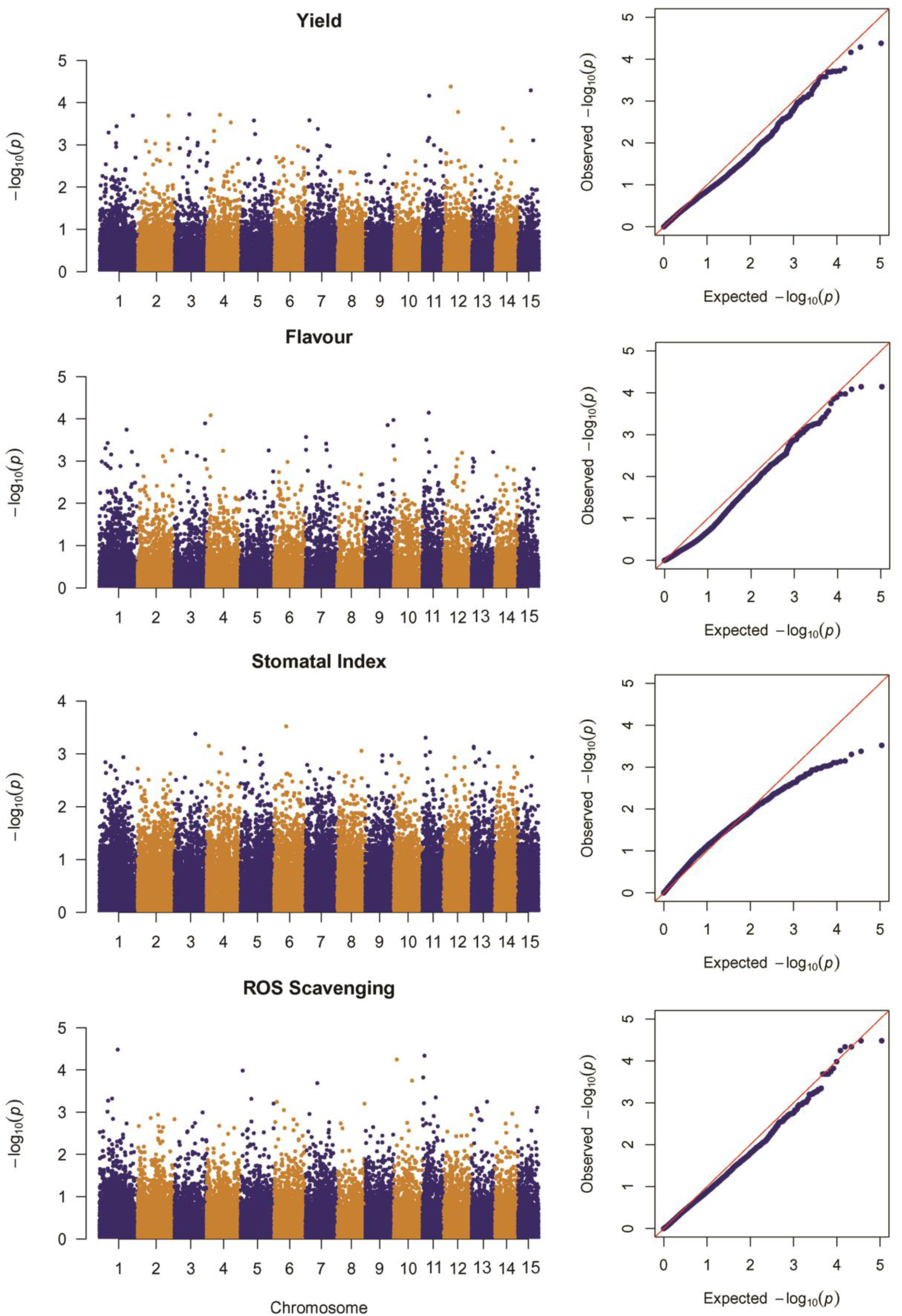
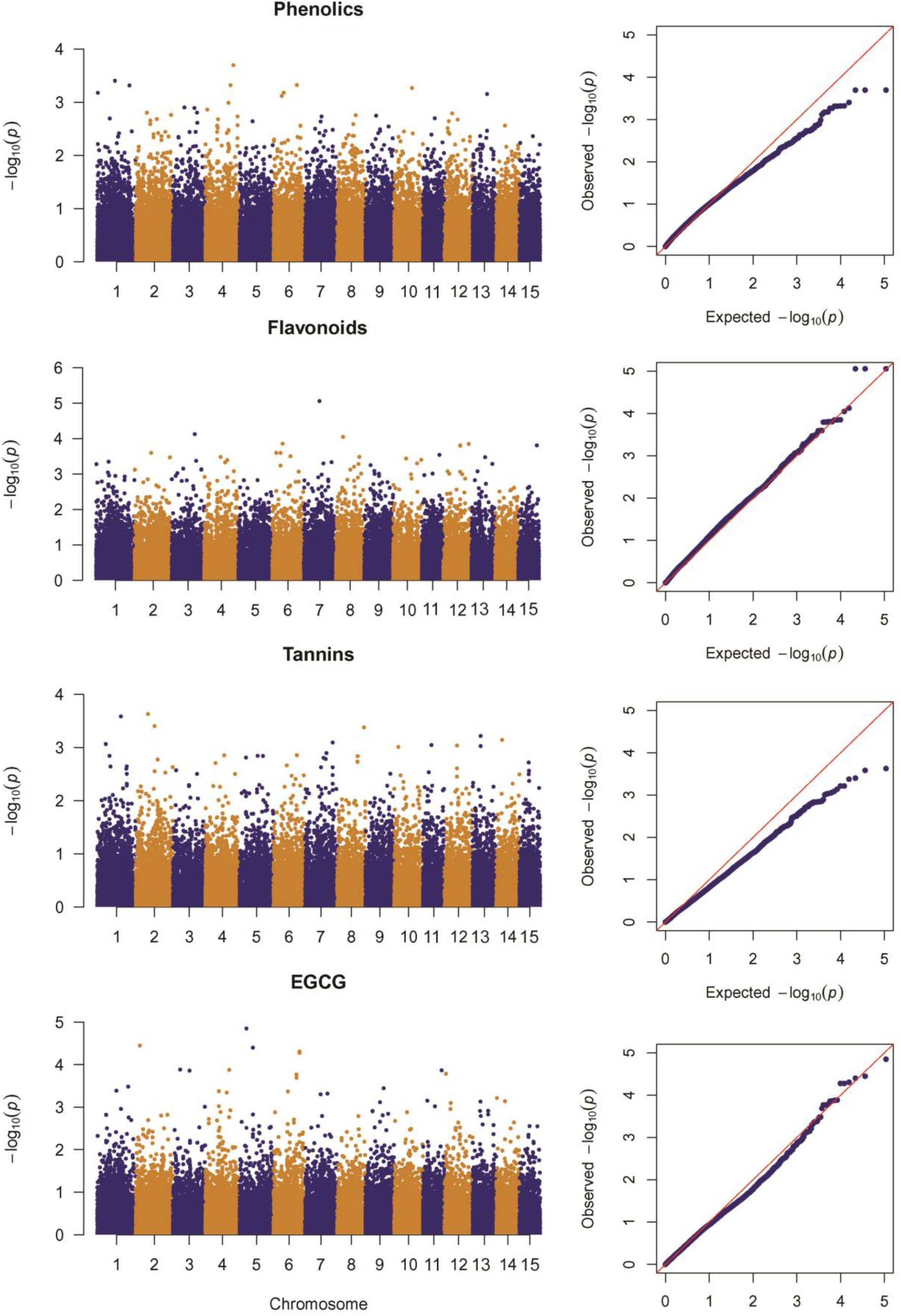
a-b - Manhattan and quantile-quantile (Q-Q) plot based on the GWAS results for eight selected traits.

## Discussion

With the advent of molecular genetic approaches, efficient genotyping with high throughput markers has led to an in-depth understanding of crop genetic resources, their management, and utilization in breeding programs (Basak et al., 2019; Sudan et al., 2019). Development of SNP markers along with the determination of their genomic position assisted the trait association analyses in plants and their diversity at an individual, community as well as species level (Guajardo et al., 2020; Huq et al., 2016; Luo et al., 2019; Mammadov et al., 2012). In the present study, ddRAD based genotyping by sequencing approach was adopted for genome-wide SNP discovery and their diversity assessment in 23 Darjeeling tea cultivars. These accessions represent contrasting genetic backgrounds, most of which are cultivated in the terrain of Darjeeling hills for their preferential quality traits. The double-digest strategy and deep coverage paired-end sequencing approach have recommended toward accomplishing greater quality SNPs, lower amounts of missing data, and precise mapping onto the reference genome (Guajardo et al., 2020; Shirasawa et al., 2016). Based on the 54K high-quality SNPs developed in this study and mapped across 15 chromosomes of tea, the genetic relatedness of the Darjeeling tea cultivars has been established. The degree of relatedness inferred through Bayesian clustering, UPGMA method, and PCoA analyses conform to each other. However, the clustering does not result in distinct lineages since the accessions carry an enormous gene pool of highly variable cultivars of Indian hybrid tea. Most of the commercial accessions considered in the present study belong to a group of ‘mixed’ ancestry as revealed by the population genetic structure of Indian tea (Raina et al., 2012). Optimum number of K obtained here for K = 2, still, three subpopulations have been assumed to concealed within the accessions as three separate origins of domesticated tea lineages exists (Meegahakumbura et al., 2018; Yamashita et al., 2019). In STRUCTURE analyses of selected Darjeeling tea cultivars, homogeneity was observed in a major part when two subpopulations were assumed. However, the same members had shown admixture between two different groups at varying degrees when K = 3 was considered. Both analyses revealed the genetic background of TS-569 cultivar to be admixed between two separate ancestries, which is following its true biparental origin (AV-2 × T-78) (Hazra et al., 2020b). The high level of heterozygosity indices (15-30%) of the studied accessions observed in the present study corroborates their ‘hybrid’ structures too. Following ddRAD based study on Chinese tea, Yang et al. (2016) also opined that due to self-incompatibility, domestication via hybridization, and climatic selection, cultivars might have resulted in broader genetic variation. A large number of high-resolution SNPs spanning tea genome thereby discovered exclusively from Darjeeling tea, which not only can distinguish the accessions with the desirable trait, but also support for understanding the effect of these variations through functional genomics aspects. Since genomic variations in the worldwide germplasm of tea is still under discovery, therefore, variants identified from this study would serve the reference collection.

Non-synonymous single nucleotide variations with the potential to alter the associated gene functions are valued in molecular breeding studies for their profound role in trait relations and marker-assisted selection (Cao et al., 2016; Guajardo et al., 2020; Shirasawa et al., 2013). Although a major fraction of SNPs discovered in this study, found to be located in the intergenic region, a substantial amount occurred in the genic region too. Among which SNPs detected in the non-coding regulatory regions such as upstream, downstream, UTR, or even intron regions might affect the gene function at the transcriptional and/or translational level. These changes along with the exon level variants should be validated on a case-by-case basis because not all SNPs in the genic or associated region are functionally impactful (Cingolani et al., 2012). The present study reveals that the occurrences of SNPs imposing modifier effects were most abundant. Similar observations were reported earlier, experimented with crops like soybean (Ramakrishna et al., 2018), pear (Montanari et al., 2019), and Prunus (Guajardo et al., 2020). Here, based on functionality, only a set of 95 genes among the total 3177 SNP impacted genes in the tea genome, could be affected highly. Because SNPs in these genes predicted to cause either stop codon gain/loss or frame-shift, missense, splice donor, and acceptor variants (Table S4). Functional annotations of high impact SNP containing genes disclosed a large proportion of them with unknown or poorly characterized function. However, the rest of them annotated to perform functions including signal transduction, secondary metabolite biosynthesis, transcriptional and translational regulation, cell cycle control, and cellular transport. Nevertheless, SNPs residing in candidate genes need further functional genomics validation to examine their impact on the consequent phenotypic effect.

Genome-wide association study has become a prospective approach for the genetic dissection of complex traits and it has successfully been adopted in various crops (Jaiswal et al., 2019; Luo et al., 2019; Raggi et al., 2019; Sudan et al., 2019). However, complex phenotypic traits linked molecular markers are still inadequate in tea to be utilized for marker-assisted selection (Hazra et al., 2019). To identify whether any nucleotide substitutions observed in the scope of present work underlying a phenotyping change, GWAS has been considered. It reveals trait-associated markers are distributed in most of the chromosomes revealing the significant role of each of the chromosomes in controlling the studied traits. Although, a large number of variants showed trait relations at a low confidence level (p < 0.05), only 1% or less retained at a more restricted parameter (p < 0.001) for most of the traits. In this study, a low sample size may stand as a limiting factor instigating a reduced significance level for the individual marker-trait association. Finally, the Bonferroni correction endorsed hardly a few loci (57 out of 54,206) to cross the threshold. The Bonferroni correction method, which estimates a threshold *p*-value to control the false positive rate, is based on the assumption that every genetic variant tested is independent of the rest (Kaler and Purcell, 2019). However, this is also considered as the most conservative method and many important loci may not qualify the stringent criterion of significance test (Wang et al., 2016). Hence, each of the markers identified can be utilized later on, for a larger population (especially from a mapping population) to check their true associations and heritability.

Since most agronomic traits are polygenic, background genetic variation of such complex traits exists in multiple loci. Genic and intergenic SNPs collectively perhaps underlie the traits and thus, understanding their functional implications is important prior to employ them in breeding studies. Exploration of the SNP surrounding genomic regions allowed the identification of several candidate genes that might have a vital role in controlling the traits. One missense variant in CSS0000649 and another variation within the upstream region of CSS0032954 exhibited significant association with flavonoid accumulation of fresh tea leaves. Functional prediction revealed the involvement of these genes in the signal transduction mechanism. Likewise, functionally relevant genic SNPs, reported in this study, might be validated and surveyed on an extensive worldwide collection of wild and cultivated genotypes of tea, so that their genealogy through domestication or artificial selection can be traced.

## Acknowledgment

Authors are thankful to the respective authority of Darjeeling Tea Research and Development Center, Kurseong, India for providing necessary permissions regarding fieldwork and to use elite accessions of Darjeeling tea for research purposes.

## Author Contribution

Conceptualization: AH and SD, fieldwork: AH and RK, laboratory experiments: AH, data interpretation: AH, CS and SD, manuscript preparation: AH and SD

## Figure and table legends

**Figure S1.**
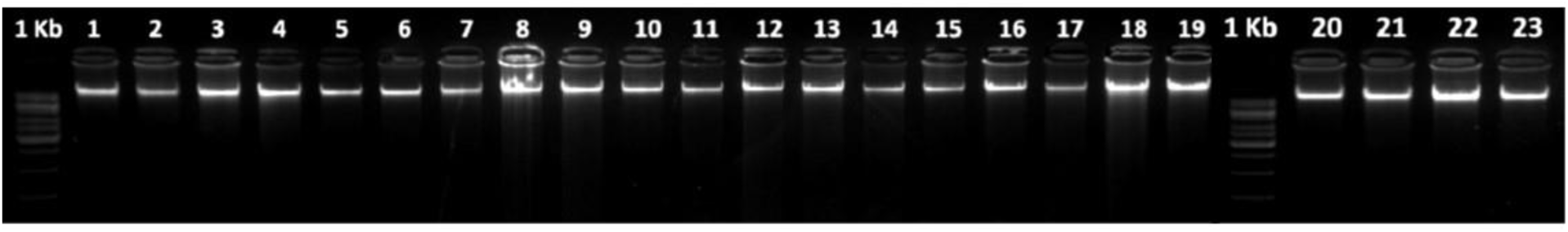
Agarose gel electrophoresis of isolated genomic DNA from 23 accessions (M – 1kb ladder)

Table S1 – List of accessions selected for the study, statistics of DNA, sequence quality, and alignment rates.

Table S2 – Summary of generated reads and variant calling

Table S3 – Records of SNP effects and putative impacts on the tea genome predicted by the SNPeff program.

Table S4 – Variant containing genes and summary of predicted effects.

Table S5 – Classification of SNP harboring genes based on KOG annotations

Table S6 - Substitution statistics of amino acid residues caused by SNPs.

Table S7 – Amount of trait-associated loci obtained for each trait with varying levels of significance.

## Notes

### Competing Interest Statement

The authors have declared no competing interest.

